# Ceasing the use of the highest priority critically important antimicrobials does not adversely affect production, health or welfare parameters in dairy cows

**DOI:** 10.1101/186973

**Authors:** Andrea Turner, David Tisdall, David C. Barrett, Sarah Wood, Andrew Dowsey, Kristen K. Reyher

## Abstract

Due to scientific, public and political concern regarding antimicrobial resistance (AMR), several EU countries have already taken steps to reduce antimicrobial (AM) usage in production animal medicine, particularly that of the highest priority critically important AMs (HP-CIAs). While veterinarians are aware of issues surrounding AMR, barriers to change such as concerns of reduced animal health, welfare or production may inhibit AM prescribing changes.

Farmers from seven dairy farms in South West England engaged in changing AM use through an active process of education and herd health planning meetings. Prescribing data was collected from veterinary sales records; production and health data were accessed via milk recording and farm-recorded data.

This study demonstrates that cattle health and welfare - as measured by production parameters, fertility, udder health, mobility data and culling rates - can be maintained and even improved alongside a complete cessation in the use of HP-CIAs as well as an overall reduction of AM use on dairy farms.

This study also identified a need to consider different metrics when analysing AM use data, including dose-based metrics as well as those of total quantities to allow better representation of the direction and magnitude of changes in AM use.

## Introduction

Antimicrobial resistance (AMR) within production animal populations is of increasing scientific, public and political concern. A recent review of published literature highlighted that, for some bacteria, AMR patterns seen in humans could be a result of antimicrobial (AM) use in livestock (1). There are also increasing reports of resistant bacteria being recovered from production animals (2-5) and evidence that the use of certain classes of antibiotics in livestock increase the risks of multiresistant bacteria being present on farms using these medicines (6, 7).

Fluoroquinolones and 3rd and 4th generation cephalosporins have been identified as ‘highest priority critically important antimicrobials’ (HP-CIAs) for human medicine by national and international bodies including the World Health Organisation (WHO), World Organisation for Animal Health (OIE) and the Food and Agriculture Organisation of the United Nations (FAO). Several EU countries have already taken steps to reduce AM use in production animal medicine(8, 9) with recent focus on the reduction of HP-CIAs. Prophylactic use of AMs is also under scrutiny as this practice has been demonstrated to increase the prevalence of multidrug resistant bacteria (10). Veterinary surgeons (VSs) in the UK have a crucial role to play in the reduction of AM administration on farms: although AMs are commonly administered by farm staff (11) (12), prescription-only veterinary medicines (POM-V) such as antibiotics may only be used if prescribed by as VS, who is therefore ultimately responsible for their use. Furthermore, as a trusted source of information to farmers (12), VSs are well-placed both to implement changes in prescription practice and to educate and motivate farmers and farm staff to engage with disease prevention strategies and more responsible AM use, all of which have been identified as ‘areas of action’ to reduce AMR (13).

Whilst many VSs understand and appreciate concerns surrounding the use of AMs in production animal medicine (14), there are also other influences on prescribing practice. Practitioners often feel that they need to respond to farmer expectations of AM prescribing (15) (16) and farmers, in turn, have concerns over potential reduced animal health, welfare and production parameters, as well as financial issues associated with reducing or changing AM use on their farms (12, 17). Despite these concerns, the majority of UK dairy farmers feel that reducing AM use in their dairy herds would be beneficial and could reduce production costs and that their peers (other farmers and VSs) would think favourably of them doing so (12). For VSs working in production animal medicine, the challenge of achieving more responsible and sustainable AM use can offer an opportunity to advance dialogue with farmers about on-farm treatment protocols, employing preventive rather than reactive management of animal health and active herd health management.

The primary aim of the study was to investigate the effect of the cessation of use of HP-CIAs at farm level on associated animal health, welfare and production parameters. Other aims were to describe the reduction in AM use using appropriate metrics and to delineate the process by which prescribing practices were changed on the participating farms. The hypothesis was that cessation of the use of HP-CIAs would not be associated with deterioration of health, welfare or production on the dairy farms investigated.

## Materials and methods

Data collected for this study originated from seven dairy farms, all of which were current clients of the Langford Farm Animal Practice (FAP), North Somerset, UK. Data were collected between 1^st^ January 2010 and 31^st^ December 2015.

### Implementing prescribing changes

In 2010, VSs at the FAP began an initiative to reduce the use of HP-CIAs on all farms under the care of the practice. During 2010-11 farmers were engaged and educated about this process through farmer meetings and herd health planning visits; during 2011-12 changes to prescribing policy and on-farm use were implemented on pilot farms (Farms 2 and 3). Subsequently, changes were extended to all farms through the following years (2012-15), including one farm that joined the practice at the end of 2013 (Farm 6). Farms worked closely with different VSs from the practice during this time, but the same changes to prescribing practice were encouraged on all farms throughout this process.

On-farm treatment protocols were changed so that HP-CIAs were not kept on farms and were only prescribed by VSs after examination of an animal. Gradually the use of HP-CIAs was phased out entirely by both farmers and VSs for any empirical treatment decision. The most significant changes were: 1) from early 2010 quinolones were no longer used or prescribed by the practice; 2) 4th generation cephalosporins used in lactating cow intramammary tubes were replaced with products containing penicillin and aminoglycoside preparations; 3) 3rd generation cephalosporins, widely used as injectable preparations, were replaced with 1st generation cephalosporins or aminopenicillins. Practice policy stated that HP-CIAs were only to be considered if indicated by the results of culture and sensitivity testing.

### Herd inclusion

Herds were included in this study if they milk recorded monthly and kept sufficiently detailed fertility and disease incidence records to allow analysis of specific health parameters (see Supplementary Materials).

### Data collection

Health, fertility and milk recording data were captured in Interherd software (Pan Livestock Services Ltd, Reading UK) or bespoke spreadsheets and then either extracted directly from these or after further analysis in another software package (TotalVet, SUM-IT Computer systems Ltd, Thame, UK; Figure 1). A full explanation of the data collection process and definition of disease parameters are described in the Supplementary Materials.

**Figure 1:**
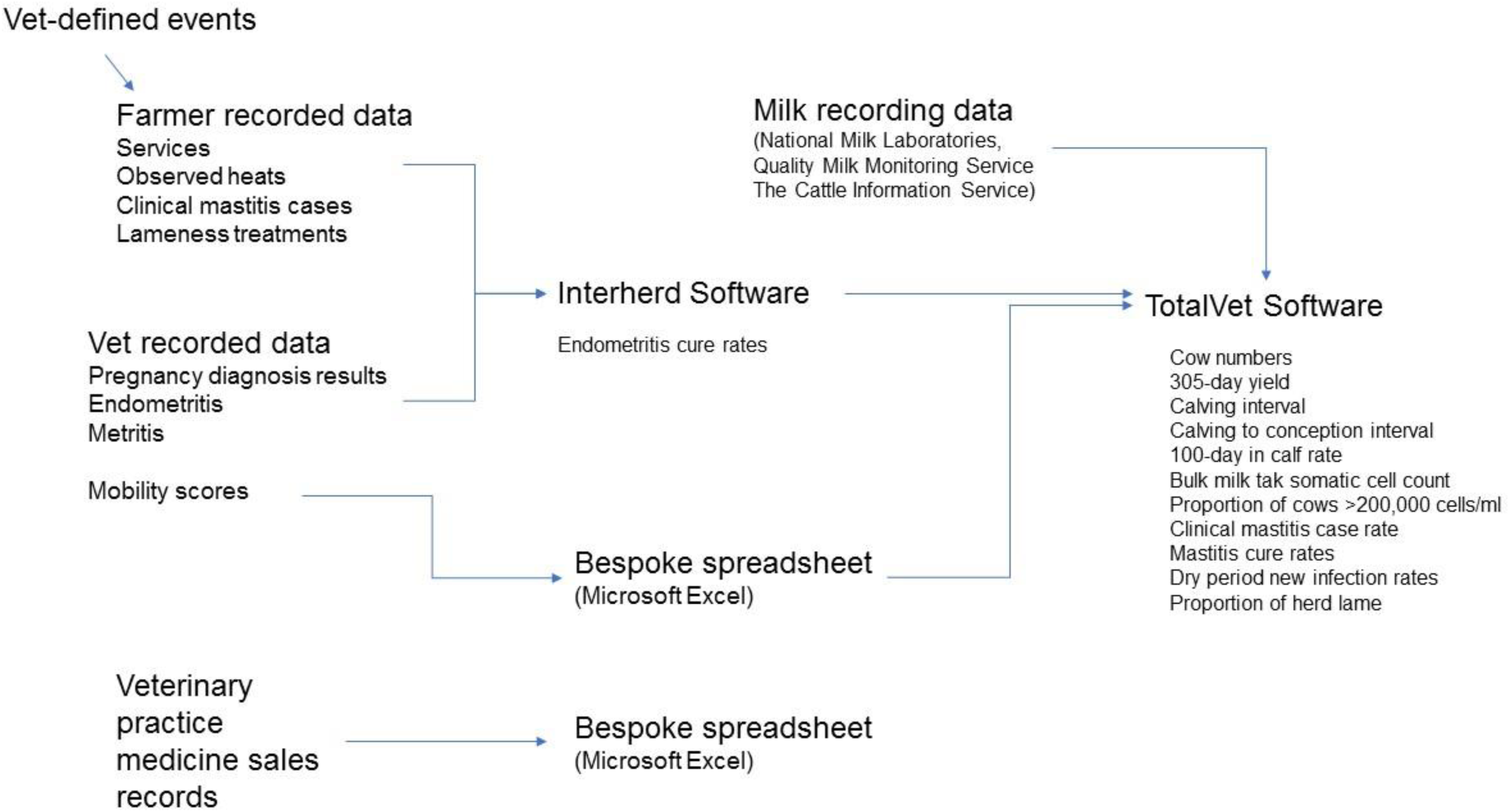
Data collection procedures used for this study and origins of calculated data parameters used in further analyses

Data for each health and welfare parameter were imported into the R programming environment, log-transformed so that regression results represented percentage change per year, and then modelled by random-intercept regression using the Bayesian MCMCglmm package(18). 130,000 MCMC samples were simulated with the first 30,000 discarded as burn-in. For each health and welfare parameter, the slope of the fitted regression curve as percentage change per year of the mean (M) and the Bayesian equivalent of confidence intervals, the 95% credible interval (CI), are reported.

### Antimicrobial prescribing

#### Data collection

Antimicrobial prescribing and sales data were accessed via practice management software (RXWorks, Newbury, UK). From this software, a list of all AM sales to each of the farms in the study was obtained for each of the study years (1st January – 31st December 2010-15). Sales data were entered into a spreadsheet (Microsoft Excel, 2013) for analysis and total kilograms (kg) of AMs used per year as well as Animal Daily Dose per animal per year (ADD), as defined by Dupont and colleagues (19), were calculated. Further information regarding the calculation of the AM use metrics and AM classification are included in the Supplementary Materials.

## Results

### Farm descriptions

All seven farms were located within a 20-mile radius of Langford FAP in South West England. All farms reared their own heifer replacements, had all-year-round calving patterns and, when housed, kept cows in freestalls. All but two farms performed selective dry cow therapy (SDCT) at the start of the study period (20) (see Supplementary Materials for SDCT protocol). The farms practicing SDCT treated, on average, 36% of cows with AMs. Mean percentage of cows dried off using AMs across all study farms across all years was 53%. Farm 6 began SDCT during 2014. Descriptive information regarding farm management and on-farm data recording are presented (Tables 1 and 2).

**Table 1.**
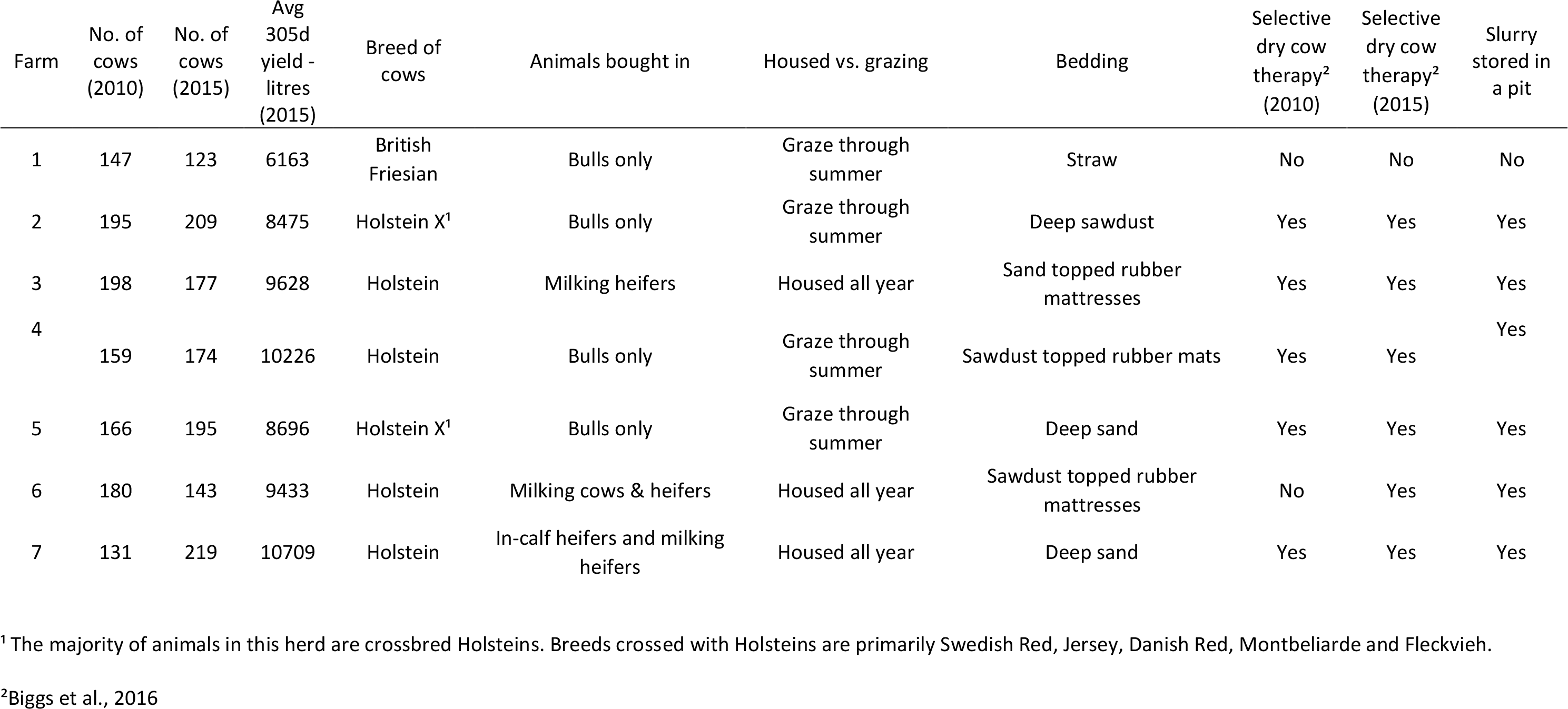
Herd information and management practices for seven dairy farms in South West England

**Table 2.** Assessment of whether reliable or accessible on-farm recorded data were available for collection and analysis from each of seven dairy farms

| Farm | Services | Observed heats | Clinical mastitis | Lameness treatments | Dry off dates | Pregnancy diagnosis results | Endo- metritis | Metritis | Mobility scores |
| --- | --- | --- | --- | --- | --- | --- | --- | --- | --- |
| 1 | Yes | No | Yes | Yes | Yes | Yes | Yes | Yes | Yes |
| 2 | Yes | Yes | Yes | Yes | Yes | Yes | Yes | Yes | Yes |
| 3 | Yes | Yes | Yes | Yes | Yes | Yes | Yes | Yes | Yes |
| 4 | Yes | Yes | Yes | No | Yes | Yes | Yes | Yes | Yes |
| 5 | Yes | No | No | No | Yes | Yes | No | No | Yes |
| 6 | Yes | Yes | Yes | Yes | Yes | Yes | Yes | Yes | No |
| 7 | Yes | Yes | Yes | Yes | Yes | Yes | Yes | Yes | Yes |

### Antimicrobial prescription

In 2010, 3^rd^ and 4^th^ generation cephalosporins were being prescribed to all seven farms (Figure 2). None of the six FAP clients were prescribed any fluoroquinolones throughout the study, meaning the reductions seen in HP-CIA use on these farms is solely due to reduced prescriptions of 3^rd^ and 4^th^ generation cephalosporins. Notably, Farm 6 (which became a client of the practice at the end of 2013) was being prescribed a similar amount of HP-CIAs from 2010-13 when compared to farms that were clients of the FAP at this time (Figure 2). Unlike the other farms, Farm 6 was prescribed small quantities of fluoroquinolones in 2011 (2.5 ADD) and 2013 (7 ADD).

**Figure 2:**
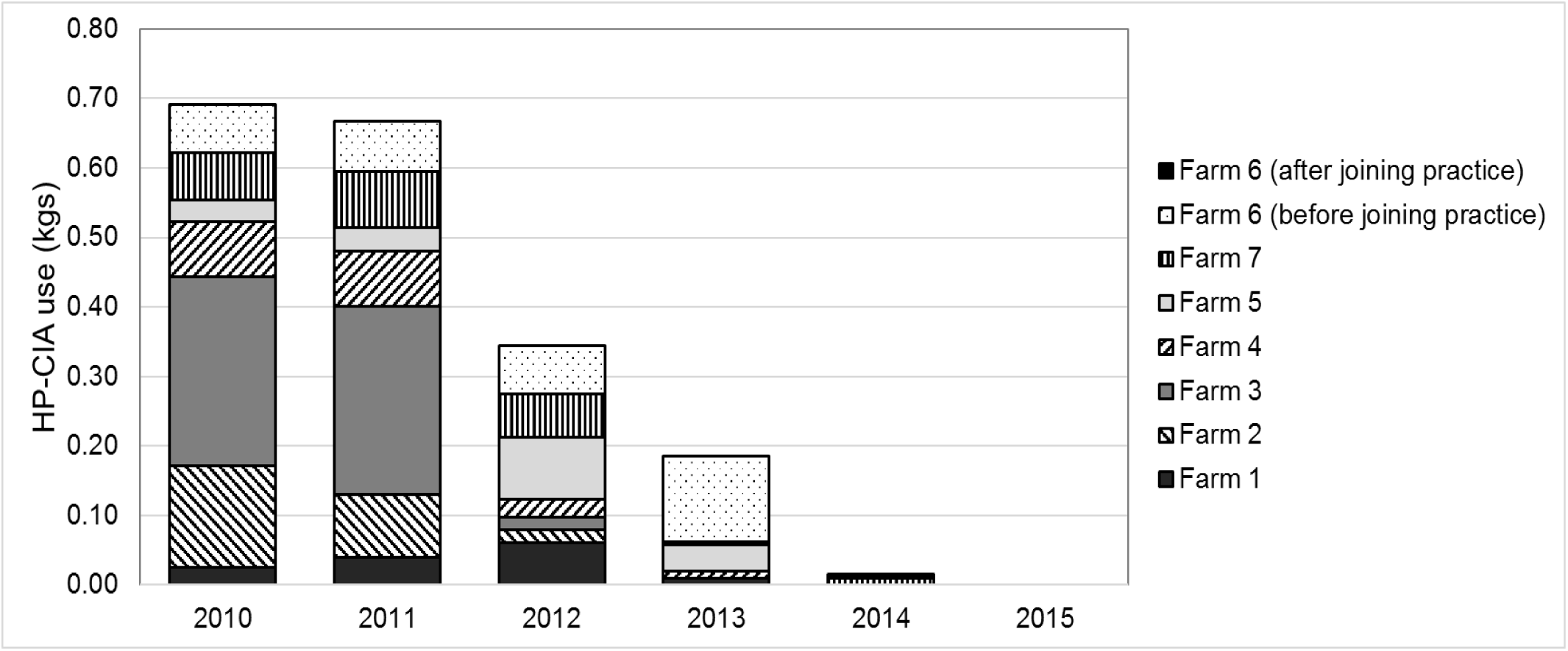
Use of highest priority critically important antimicrobials across all seven study farms, in total kilograms (kg), by year. Farm 6 joined the Langford Farm Animal Practice at the end of 2013.

Due to the changes in prescribing practices, the total quantity of HP-CIAs reduced from 0.9 kg or 7.2 ADD in 2010 (accounting for 6.3% and 41.0% of kg and ADD, respectively) until they were not prescribed to any farms in 2015. HP-CIA prescriptions consistently accounted for a larger proportion of ADD than kg of AMs.

The majority of HP-CIAs prescribed in 2010 across the seven farms were licenced for systemic administration (77% kg, 61% ADD; Table 3). Intramammary (lactating cow) preparations containing 3^rd^ and 4^th^ generation cephalosporins also significantly contributed to total quantities of HP-CIAs prescribed (23% kg, 39% ADD), although no dry cow intramammary preparations containing HP-CIAs were prescribed throughout the six years of the study. From 2010 to 2015, HP-CIA-containing intramammary tubes constituted 26, 24, 8, 3, 0 and 0% of all intramammary preparations, respectively, by kg and 57, 58, 21, 8, 0 and 0%, respectively, by ADD.

**Table 3:** Licenced routes of administration of antimicrobials prescribed to seven study farms each year from 2010 to 2015 calculated as total kilograms (kg) and animal daily doses (ADDs). Antimicrobials are classed as first line or highest priority critically important antimicrobials (HP-CIAs); intramammary tubes are split by lactating cow (LC) and dry cow (DC) treatments.

|  | 2010 |  | 2011 |  | 2012 |  | 2013 |  | 2014 |  | 2015 |  |
| --- | --- | --- | --- | --- | --- | --- | --- | --- | --- | --- | --- | --- |
|  | kg | ADD | kg | ADD | kg | ADD | kg | ADD | kg | ADD | kg | ADD |
| Systemic 1st line | 10.8 | 6.5 | 8.3 | 7.0 | 13.4 | 17.1 | 15.4 | 21.1 | 13.1 | 11.7 | 11.3 | 9.7 |
| Systemic HP-CIAs | 0.7 | 4.4 | 0.1 | 4.1 | 0.4 | 2.0 | 0.1 | 1.1 | 0.0 | 0.1 | 0.0 | 0.0 |
| Intramammary LC 1st line | 0.4 | 2.2 | 0.6 | 2.1 | 0.8 | 4.0 | 0.7 | 3.4 | 1.1 | 4.5 | 0.8 | 4.8 |
| Intramammary LC HP-CIAs | 0.2 | 2.8 | 0.2 | 3.0 | 0.1 | 1.1 | 0.0 | 0.3 | 0.0 | 0.0 | 0.0 | 0.0 |
| Intramammary DC 1st line | 0.9 | 1.5 | 0.9 | 1.7 | 0.9 | 1.1 | 0.9 | 1.1 | 0.9 | 1.0 | 1.1 | 1.5 |
| Topical 1st line | 0.0 | 0.1 | 0.0 | 0.1 | 0.0 | 0.1 | 0.0 | 0.0 | 0.0 | 0.0 | 0.0 | 0.2 |
| Intrauterine 1st line | 0.1 | 0.6 | 0.1 | 0.3 | 0.0 | 0.8 | 0.0 | 0.3 | 0.0 | 0.6 | 0.0 | 0.6 |
| TOTAL | 13.0 | 18.1 | 10.2 | 18.3 | 15.5 | 26.1 | 17.1 | 27.4 | 15.1 | 17.9 | 13.2 | 16.7 |

Overall, systemic AMs accounted for the largest quantities of AM prescriptions to each farm (by both kg and ADD) followed by intramammary preparations. The use of intrauterine and topical AMs was low across all years (Table 3).

Both metrics show variation in the quantities of AMs being prescribed to the study farms each year; both the total kg of AMs prescribed and the total number of ADD prescribed to the study farms increased in 2012 and 2013 before reducing through 2014 and 2015. When represented in ADD, AM prescription in 2015 was lower than in any previous year of the study (16.7 ADD; Table 3).

The mean AM prescription across all study farms was also calculated in mg/kg for each year of the study and was found to be 14.0, 12.0, 17.5, 18.0, 15.1 and 12.7 mg/kg respectively between 2010 and 2015. The minimum value at the farm level was 5.0 mg/kg and the maximum value was 35.3 mg/kg for individual farms across all years of the study.

The increase in the total quantity of AMs prescribed in 2012, as measured in both kg and ADD, was largely due to an increase of penicillins (including penicillin in combination with streptomycin), and 1st generation cephalosporins (Table 4). There was little change in the prescription of anti-pneumonial AMs considered to be used primarily for the treatment of calves (tetracyclines, amphenicols and macrolides, including long-acting macrolides) over the six years of the study (2.2, 2.0, 1.7, 2.3, 2.1 and 1.8 kg, respectively, through 2010-15), accounting for 0.1 ADD in 2010 and decreasing to 0.08 ADD in 2015.

**Table 4:** Classes of antimicrobials prescribed to seven study farms in 2010 – 2015, in kilograms (kg) and animal daily doses (ADDs)

|  | 2010 |  | 2011 |  | 2012 |  | 2013 |  | 2014 |  | 2015 |  |
| --- | --- | --- | --- | --- | --- | --- | --- | --- | --- | --- | --- | --- |
|  | kg | ADD | kg | ADD | kg | ADD | kg | ADD | kg | ADD | kg | ADD |
| 3rd generation cephalosporins | 0.5 | 3.3 | 0.4 | 3.2 | 0.2 | 1.5 | 0.1 | 1.0 | 0.0 | 0.1 | 0.0 | 0.0 |
| 4th generation cephalosporins | 0.2 | 3.8 | 0.2 | 3.9 | 0.1 | 2.1 | 0.0 | 0.5 | 0.0 | 0.0 | 0.0 | 0.0 |
| Fluroquinolones | 0.0 | 0.0 | 0.0 | 0.0 | 0.0 | 0.0 | 0.0 | 0.1 | 0.0 | 0.0 | 0.0 | 0.0 |
| Penicillins | 4.2 | 4.8 | 4.0 | 5.0 | 6.3 | 7.3 | 7.2 | 7.1 | 5.8 | 5.2 | 5.5 | 5.0 |
| Amoxicillins with clavulanic acid | 0.0 | 0.9 | 0.0 | 0.6 | 0.3 | 7.4 | 0.4 | 9.5 | 0.3 | 6.1 | 0.4 | 6.2 |
| 1st generation cephalosporins | 0.5 | 1.5 | 0.3 | 1.2 | 1.6 | 2.3 | 1.4 | 2.2 | 2.1 | 3.0 | 1.6 | 2.3 |
| Potentiated sulphonamides | 0.2 | 1.6 | 0.4 | 1.0 | 0.5 | 0.7 | 1.3 | 1.2 | 1.1 | 0.9 | 0.5 | 1.2 |
| Macrolides | 0.0 | 0.2 | 0.0 | 0.4 | 0.1 | 0.6 | 0.0 | 1.2 | 0.1 | 0.5 | 0.0 | 0.2 |
| Lincosamides | 0.0 | 0.0 | 0.0 | 0.0 | 0.0 | 0.2 | 0.0 | 0.0 | 0.0 | 0.1 | 0.0 | 0.0 |
| Chloramphenicol derivatives | 0.9 | 0.0 | 0.5 | 0.0 | 0.4 | 0.0 | 0.5 | 0.0 | 0.2 | 0.0 | 0.1 | 0.0 |
| Aminoglycosides | 4.2 | 0.2 | 2.8 | 0.1 | 4.6 | 0.1 | 4.1 | 0.1 | 3.5 | 0.0 | 3.3 | 0.0 |
| Aminocoumarins | 0.1 | 0.2 | 0.1 | 1.8 | 0.2 | 3.4 | 0.2 | 3.2 | 0.2 | 1.1 | 0.1 | 0.6 |
| Tetracyclines | 2.2 | 1.6 | 1.5 | 1.0 | 1.2 | 0.7 | 1.9 | 1.2 | 1.8 | 0.9 | 1.8 | 1.2 |
| TOTAL | 13.0 | 18.1 | 10.2 | 18.3 | 15.5 | 26.1 | 17.1 | 27.4 | 15.1 | 17.9 | 13.2 | 16.7 |

### Production parameters

Mean 305-day milk yield showed an increasing trend, with a mean increase of 0.6% per year over the six years of the study (95% CI [-0.2, 1.4]; Figure 3).

**Figure 3:**
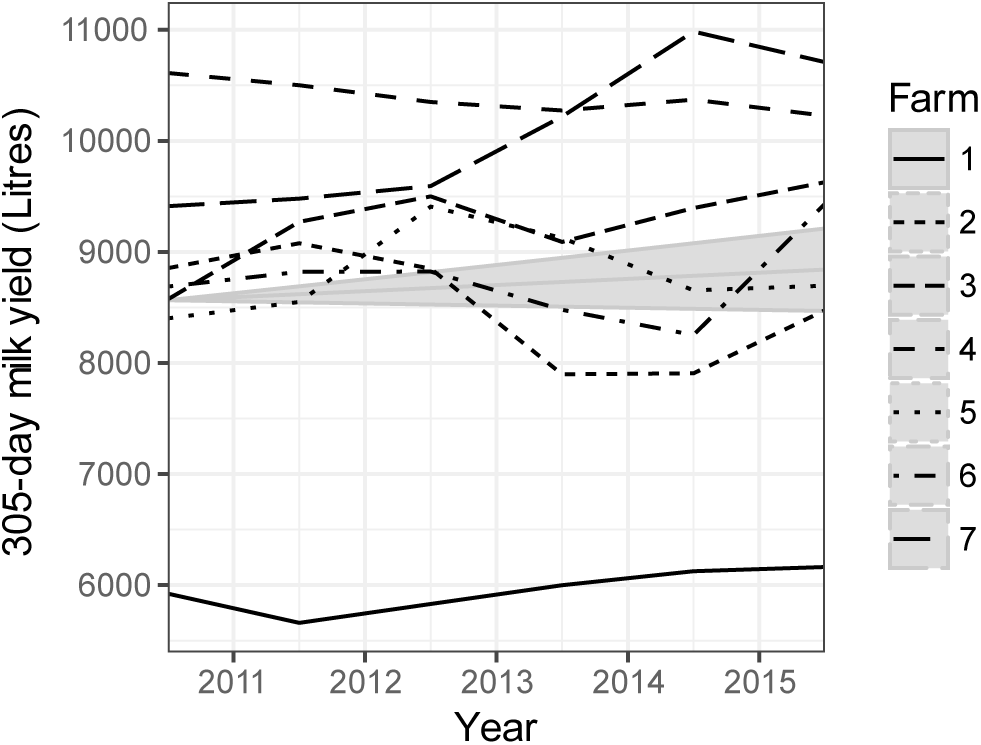
Mean 305-day yield from all seven farms. Grey areas show the variation in percentage change per year as compared to the mean (M, central grey line), and the intercept on the y-axis represents the geometric mean value across seven farms in 2010. Top and bottom grey lines represent upper and lower 95% credible intervals (CI), respectively. M = 0.6, CI = [-0.2, 1.4].

### Fertility parameters

Fertility parameters showed an improving trend over the course of the six-year study period (Figure 4). Calving index decreased from 413 days in 2010 to 390 days in 2015 (M −0.9 %; 95% CI []-1.4, −0.5). Mean 100-day in-calf rate increased across the seven farms from 0.4 in 2010 to 0.5 in 2015 (M 5.0%; 95% CI [1.8, 7.8]). During the same period, mean calving to conception interval showed a decreasing trend from 147 days in 2010 to 120 days in 2015 (M −2.8%/yr; 95% CI −5.8, 0.2). During the study period, the mean endometritis failure to cure rate is unlikely to have changed (M −5.7%/yr; 95% CI [-19.5, 7.6]).

**Figure 4:**
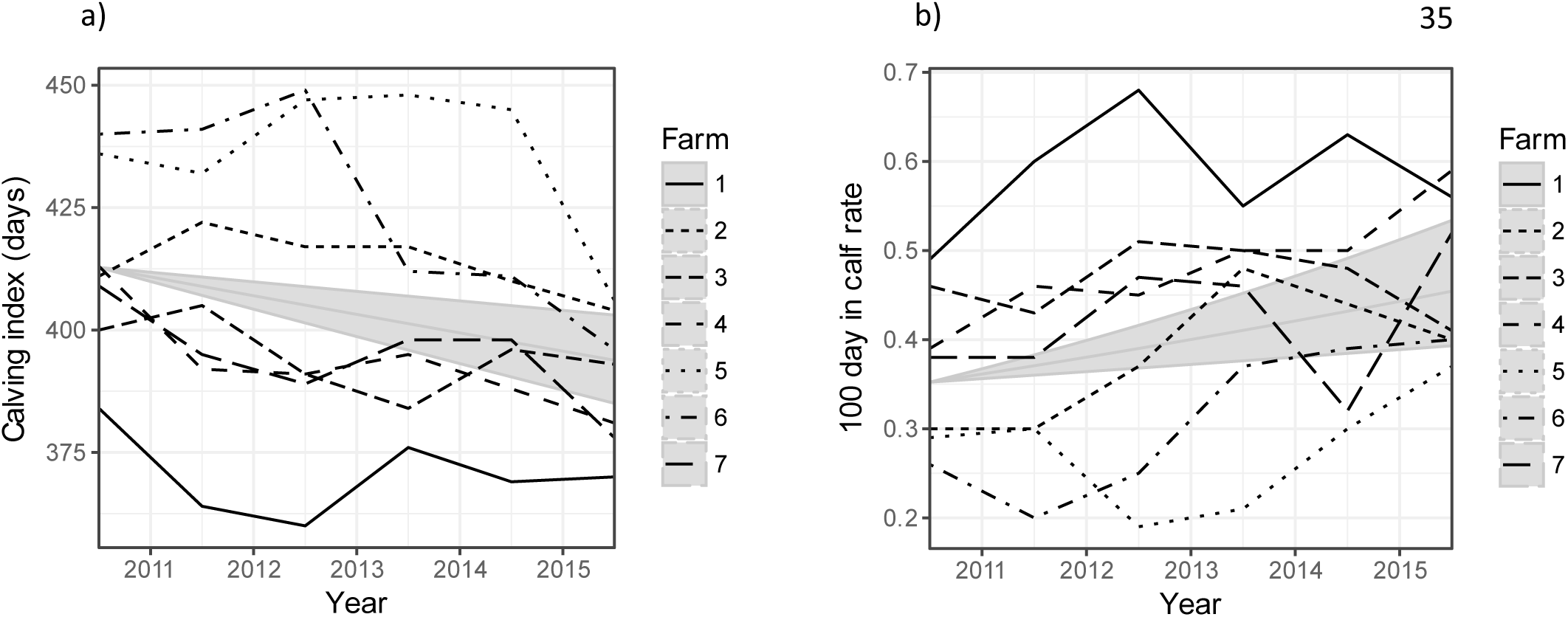
a) Calving index and b) 100-day in-calf rate for each of seven farms across six years. Grey areas show the variation in percentage change per year as compared to the mean (M, central grey line), and the intercept on the y-axis represents the geometric mean value across seven farms in 2010. Top and bottom grey lines represent upper and lower 95% credible intervals (CI), respectively. Calving index M = −0.9, 95% CI −1.4, −0.5; 100-day in-calf rate M = 5.0; 95% CI [1.8, 7.8].

### Udder health parameters

Clinical mastitis cases (as recorded by farmers) decreased on five of the seven farms between 2010 and 2015 (Figure 5A). On Farm 1, clinical mastitis cases rose from 10 cases/100 cows/year to 29 cases/100 cows/year. The on-farm records of clinical mastitis for Farm 5 were not complete and so are not included in the analysis. From 2010-15, mean mastitis cure rates (as measured by somatic cell counts, SCC) showed a mean increase of 7.5%/yr (95% CI [1.2, 13.3]; Figure 5B). Mean dry cow cure rates showed a decreasing trend (M −1.3%; 95%CI [−2.9, 0.2]).

**Figure 5:**
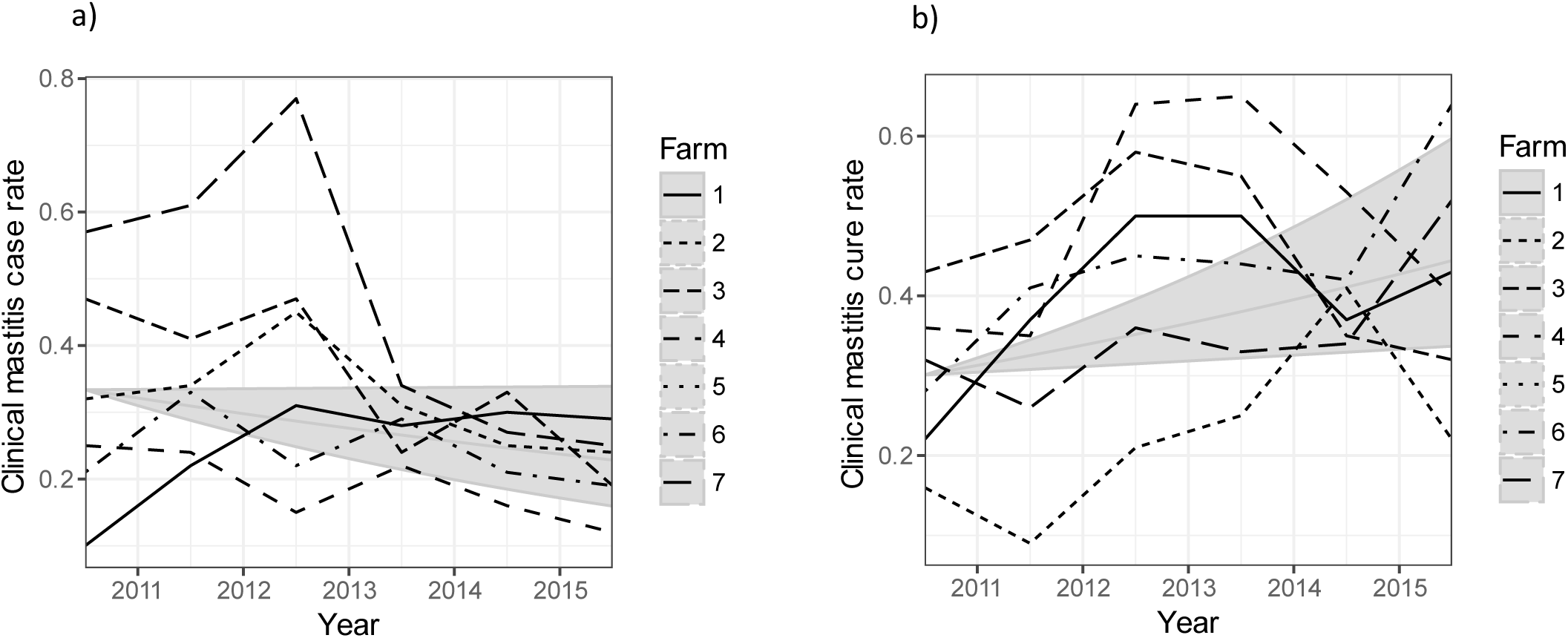
a) Clinical mastitis case rates per cow per year; b) Clinical mastitis all case cure rates per cow per year (n.b. Farm 5 is not represented as on-farm clinical mastitis records were not complete). Grey areas show the variation in percentage change per year as compared to the mean (M, central grey line), and the intercept on the y-axis represents the geometric mean value across seven farms in 2010. Top and bottom grey lines represent upper and lower 95% credible intervals (CI), respectively. Clinical mastitis case rate M = −7.8, 95% CI [-16.1, 0.1]; Clinical mastitis cure rates M = 7.5, 95% CI [1.2, 13.3].

Subclinical mastitis risks, as assessed by mean proportion of cows with SCC over 200,000 cells/ml and the mean percentage of cows chronically infected (SCC>200k for two consecutive milk recordings) are unlikely to have changed (M 1.0%; 95% CI [−2.2, 4.4] and M 2.2%, 95% CI [−2.1, 6.9], respectively). Variation in the 12-month rolling bulk milk SCC which ranged from 161 in 2013 to 262 in 2014 is largely due to extreme variability in the bulk milk SCC on one farm (Farm 1; Figure 6).

**Figure 6:**
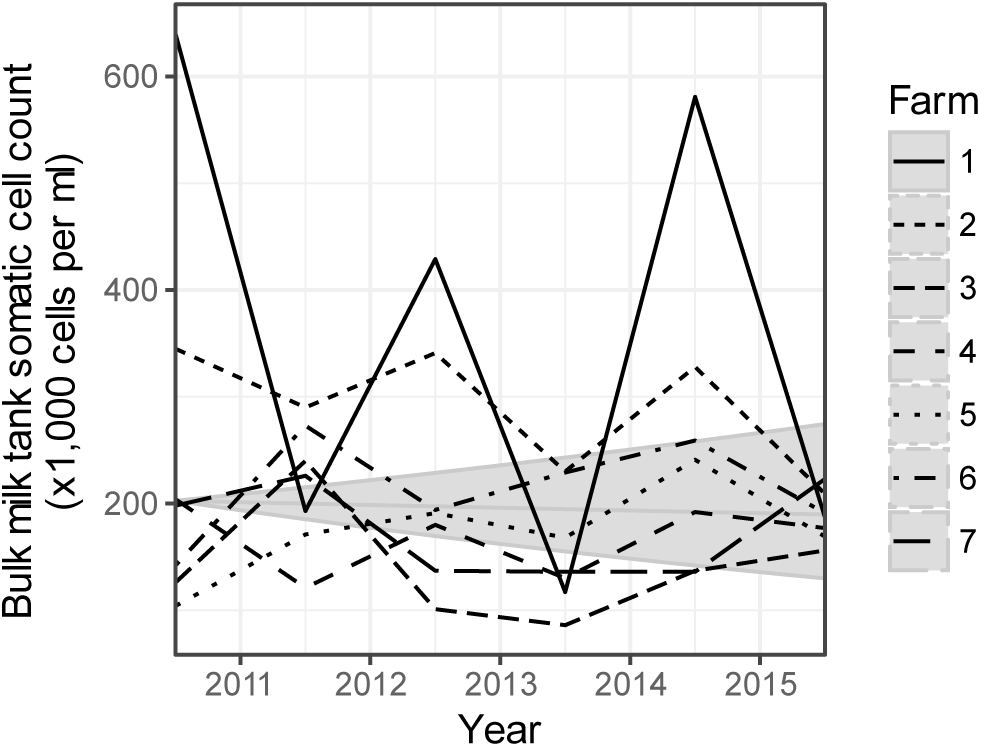
Geometric mean of the bulk milk tank somatic cell count (BMTSCC) for each of seven farms during each year of the study. Grey areas show the variation in percentage change per year as compared to the mean (M, central grey line), and the intercept on the y-axis represents the geometric mean value across seven farms in 2010. Top and bottom grey lines represent upper and lower 95% credible intervals (CI), respectively. BTMSCC M = − 1.4, 95% CI [−9.4, 5.9].

### Lameness

The percentage of cows scored as lame at mobility scoring sessions performed by VSs decreased over the six years of the study on all farms with complete data (Figure 7).

**Figure 7:**
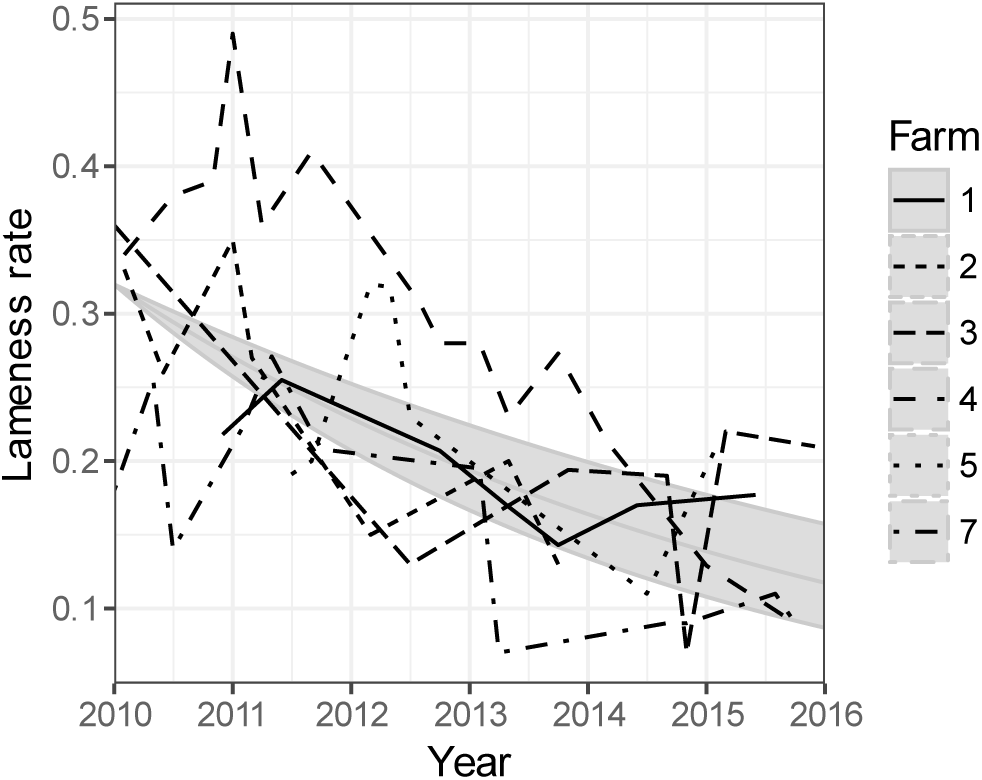
Lameness rates (cases per cow per year) at mobility scoring performed on six farms by veterinary surgeons between 2010 and 2015. Y-axis shows the lameness rates per cow per year (n.b. Farm 6 is not represented as mobility scoring records were not complete). Grey areas show the variation in percentage change per year as compared to the mean (M, central grey line), and the intercept on the y-axis represents the geometric mean value across seven farms in 2010. Top and bottom grey lines represent upper and lower 95% credible intervals (CI), respectively. Lameness rate M = −18.2, 95% CI [−24.1, −12.5].

### Culling

Culling percentages ranged from 13 to 33% across the seven farms in 2010 and remained within this range throughout subsequent years of the study (M −3.7%; 95% CI [−7.9, 0.3]; Figure 8). While certain farms decreased the percentage of the herd culled each year (Farm 2), others increased the culling percentage overall (Farm 6), and there was variability both within and between farms.

**Figure 8:**
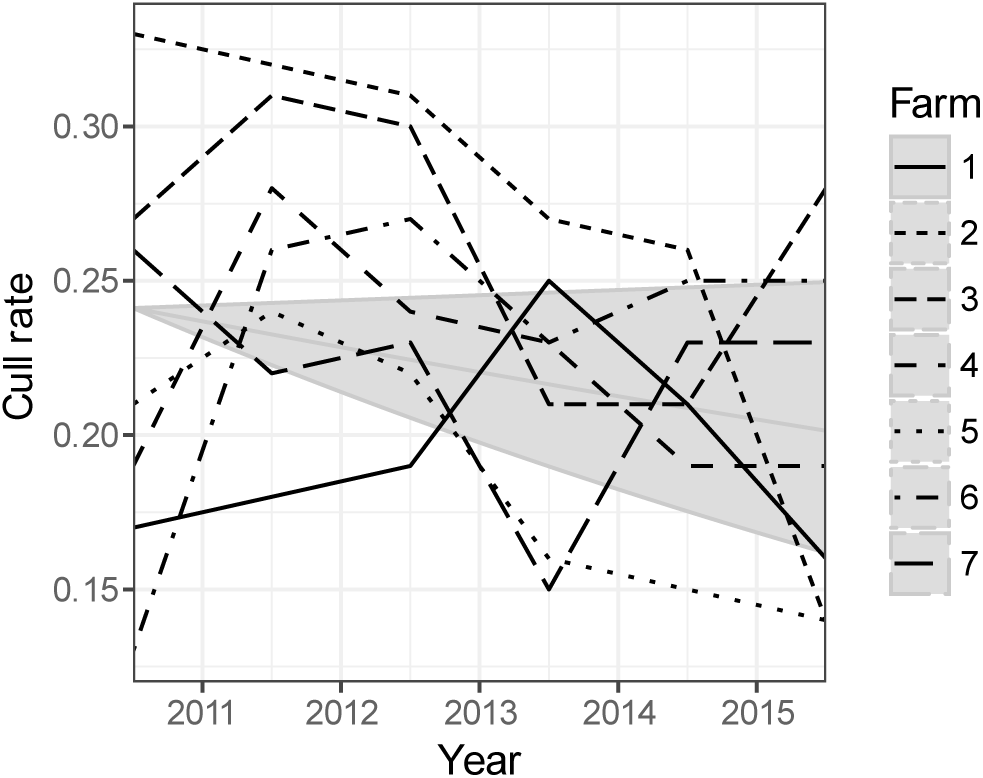
Cull rate (per cow per year). Y-axis shows the cull rate. Grey areas show the variation in percentage change per year as compared to the mean (M, central grey line), and the intercept on the y-axis represents the geometric mean value across seven farms in 2010. Top and bottom grey lines represent upper and lower 95% credible intervals (CI), respectively. Cull rate M = −3.7, 95% CI [−8.1, 0.6].

## Discussion

The aims of this study were to describe how prescribing policy was changed to achieve a cessation in HP-CIA use on seven dairy farms, to appropriately assess these AM use changes using two different metrics and to examine the effects of discontinued HP-CIA use on cattle health and welfare.

Farms were selected as a convenience sample due to inclusion criteria. It is accepted that for these reasons, and as clients of a first-opinion farm animal practice connected with a veterinary school, these farmers may not be representative of all UK dairy farms. However, all the study farms were commercial dairies and the information outlined in Table 1 demonstrates that these farms represent herd sizes and management systems commonly found in the UK.

In this study, AMs prescribed to farms were assumed to represent AM use. It is possible that some prescribed AMs may not have been administered to animals, but may have gone unused or been disposed of. For this reason, AM prescription is likely to overestimate use, however the authors judged this unlikely to be significant as each farm had weekly or bi-weekly veterinary visits, making it unnecessary for AMs to be ‘stockpiled’.

### Calculation of antimicrobial use

The comparison of AM prescribing using two metrics (Tables 3 and 4) highlights the degree of change implemented by VSs and farmers; HP-CIAs represent a small proportion of the quantity (kg) of AMs used in 2010-12, but represent a far larger proportion of ADD administered in the same years. This is particularly true of the use of HP-CIAs in intramammary tubes. The way in which the ADD metric is reported alongside kg of AMs prescribed demonstrates the ways in which different analyses of AM use can represent these data differently. Calculation of mean AM consumption in mg/kg values for each year of the study was included to demonstrate the AM prescription of these herds in comparison with the 50 mg/kg target suggested by the O’Neill Report (1) and adopted for the UK across the combined livestock sectors(21). It should be noted that a move away from HP-CIAs towards other AM treatments is likely to increase the total mg/kg use due to differences in dose rates. Conversely driving down total mg/kg targets could have unintended consequences if veterinary surgeons were to increase their reliance on low dose HP-CIAs.

### Antimicrobial prescription

Similar to other studies (22, 23), penicillins and 1st generation cephalosporins were prescribed in the highest quantity (kg) in 2014 and 2015. Tetracyclines were also commonly prescribed, as has been previously reported (22). This study differs from other studies which only analysed medicines considered to be primarily used to treat the milking herd (23). These findings suggest that the groups of AMs used most frequently by farms in this study are also commonly used on other dairy farms in the developed dairy areas of the world.

The majority of AMs were administered systemically, which corresponds to the findings of Saini and colleagues (22), but was not true for all farms in other studies (24). The difference in these findings is likely to be because in both this study and the study by Saini and colleagues, medicines used for calves and youngstock were included, whereas medicines used only in adults were included by Stevens and colleagues (24).

After prescribing changes were implemented on farms during 2011-12, the total quantities of AMs prescribed in kg increased by 52% (Table 4). This was in part due to the differences in mgs/dose of first line antimicrobials and HP-CIAs resulting in the total use of AMs appearing falsely elevated, demonstrated by a smaller increase (25%) as represented by ADD. However, across both metrics, the quantities of AMs prescribed during 2012 and 2013 were higher than in 2011, suggesting increased use to treat disease in these years.

### Health and production parameters

Milk yield on these farms remained stable over the study period, demonstrating that production was maintained alongside a cessation of HP-CIA use and an overall decrease in ADD prescribed through 2014-15 (Figure 3).

Prior to prescription policy changes made in 2011-2012, farmers would frequently treat cows that were systemically unwell due to retained foetal membranes or metritis with injectable 3rd generation cephalosporins. Subsequently to prescription policy changes, cattle with these diseases were treated with aminopenicillin preparations or amoxicillin-clavulanic acid combinations. It is assumed that a significant deterioration in uterine health would have negative impacts on fertility parameters (25, 26), however no negative effects on fertility parameters were observed in association with the reduction in HP-CIAs on these farms; in fact, the calving index decreased, the 100-day in-calf rate increased and mean number of days from calving to conception showed a decreasing trend through 2010-15 (Figure 4).

The protocol for the antibiotic treatment of endometritis (intrauterine 1st generation cephalosporin licenced for this purpose in cattle) did not change on any of the farms throughout the study period. It is therefore unsurprising that the mean failure to cure rate of endometritis remained stable.

Mean clinical mastitis case rates are reported for reference (Figure 5A), and it should be noted that they are lower than the average clinical mastitis case rates on farms in England and Wales (estimated to be 47 cases/100 cows/year (equating to a case rate of 0.47) (27)). The increase in clinical mastitis case rate in 2012/13 corresponds to the increase in prescription of AMs during these years and so is likely to contribute to the increased AM use in these years. Despite variation in clinical mastitis case rates throughout the six study years, the clinical mastitis cure rates improved, suggesting that the treatments being used through 2014 and 2015 (none of which contained HP-CIAs) appeared to be at least as effective at achieving a cure as the AMs being used in 2010, when 40% of ADD of intramammary preparations prescribed contained HP-CIAs. This is consistent with a study in which implementation of a restricted AM usage policy in Dutch herds did not impact negatively on udder health (28). HP-CIAs were not used in dry cow treatments in 2010; dry cow treatment protocols were therefore not changed between 2010 and 2015. The trend towards lower dry cow cure rates may be due to changes in mastitis pathogens, changes in dry cow management or in decreased efficacy of dry cow intramammary treatments, but it is difficult to differentiate between these with the data available.

Prior to prescription policy changes, infectious causes of lameness in milking cows (in particular interdigital necrobacillosis) would commonly have been treated with injectable 3^rd^ generation cephalosporin preparations. Although primarily affected by management factors, lameness rates would be expected to have increased if infectious causes of lameness had a poor treatment recovery rate. The fact that mobility scores from all six farms for which there are data available shows a general reduction in the rates of lameness at mobility scores indicates that foot health can also be maintained alongside reduced HP-CIA use, specifically through the use of 1st generation cephalosporins and aminopenicillins in place of 3rd generation cephalosporins.

It is possible that animal health may have been deteriorating without altering the health parameters assessed if culling rates on the farms had increased significantly. Maximum culling rates through 2013-15, however, were lower than those of 2010-12, while the minimum culling rate remained at a similar level. Mean percentage change and associated CIs suggest that culling rates were maintained through all six years of the study (Figure 7).

As this was a retrospective study, only health data and production parameters that were available from milk recording data and on-farm records could be analysed. Rather than assess the efficacy of specific treatment changes for particular diseases, this study sought to provide evidence about the effects on production and welfare parameters of reducing HP-CIAs on dairy farms. Where possible, individual animal treatment outcomes were assessed and presented. Although many treatment outcomes may be of interest (including outcomes of pneumonia treatments in calves, clinical cure rates of pneumonia in calves or cases of interdigital necrobacillosis in dairy cows), not all of these could be assessed due to insufficient or inappropriate on-farm records. While calf health parameters were not directly assessed in these analyses, it should be noted that prescriptions of AMs classed as anti-pneumonials remained stable between 2010 and 2015 (both in kg and ADD). Hence it appears that calf health (specifically regarding respiratory disease as AMs were not used to treat diarrhoeal disease) did not deteriorate during this time.

Future work in this area should consider prospective evaluations of individual animal treatment outcomes for specific diseases over time as AM use changes, with consistent follow-up of cases and recording of clinical or bacteriological cures.

### Changing prescribing practice

While HP-CIAs were used on all seven farms at the beginning of the study period, some farms had higher use than others, and the time taken to implement strategies to reduce HP-CIA use also varied between farms. This may reflect, in part, both the farmers’ willingness to change AM use and individual VSs’ willingness to alter on-farm treatment protocols or their own prescribing practice. Within two years of practice-wide implementation of prescribing changes, however, all six of the seven study farms that were clients of the FAP at the start of the study period had either significantly reduced or altogether ceased use of HP-CIAs. Furthermore, the farm that joined the practice at the end of 2013 (Farm 6) changed its use of HP-CIAs with immediate effect after joining the practice and initiated use of SDCT, providing evidence that prescribing changes can be enacted quickly on farms.

The changes to prescribing practice demonstrated in this study were made alongside various strategies to improve animal management and husbandry on the farms which subsequently resulted in a net decrease in doses (ADD) of AMs being prescribed, despite an increase in the use of certain AM classes. Reducing the need to prescribe AMs by working to improve herd health rather than forcibly avoiding the prescription or use of AMs offers a sustainable way to safeguard animal health and welfare and maintain food production alongside reduced AM use.

A cessation of the use of HP-CIAs and a decrease in the use of AMs within the livestock industry should be a key target for farmers and VSs, and has been shown to be achievable whilst maintaining animal health, welfare and production.

Data and analyses of the sort presented here are crucial to addressing barriers that farmers and VSs face when deciding to alter their AM use and provide much-needed evidence to aid a shift in the ‘social norm’ of AM use in production animal medicine.

### Conclusions

This is the first study to demonstrate that cattle health and welfare - as measured by culling rates, production parameters, fertility, udder health and mobility data - can be maintained, if not improved, alongside a complete cessation in the use of HP-CIAs as well as an overall reduction of AM use on dairy farms when prescribing practices are altered in line with proactive herd health planning strategies.

## Acknowledgements

The authors wish to thank to the VSs and clients of Langford FAP for their assistance in collecting and allowing access to their farm data. Thanks also to the members of the University of Bristol’s Farm Animal Group for their input and advice, and to the Pat Impson Memorial Fund and Langford Trust for supporting the work via their sponsorship of the Pat Impson Residency in Bovine Health Management.

## Funding

This work was supported by the Pat Impson Memorial Fund and Langford Trust via their sponsorship of the Pat Impson Residency in Bovine Health Management at the University of Bristol Veterinary School.

## Transparency declarations

None to declare.

